# Solid Tumor Treatment via Augmentation of Bioactive C6 Ceramide Levels with Thermally Ablative Focused Ultrasound

**DOI:** 10.1101/2023.03.23.532394

**Authors:** E. Andrew Thim, Todd Fox, Tye Deering, Luke R. Vass, Natasha D. Sheybani, Mark Kester, Richard J. Price

**Author notes:** Corresponding Author: Richard J. Price, Ph.D., Department of Biomedical Engineering, Box 800759, Health System, University of Virginia Charlottesville, VA 22908, USA.

## Abstract

Sparse scan partial thermal ablation (TA) with focused ultrasound (FUS) may be deployed to treat solid tumors and increase delivery of systemically administered therapeutics. Further, C6-ceramide-loaded nanoliposomes (CNLs), which rely upon the enhanced permeation and retention (EPR) effect for delivery, have shown promise for treating solid tumors and are being tested in clinical trials. Here, our objective was to determine whether CNLs synergize with TA in the control of 4T1 breast tumors. CNL-monotherapy of 4T1 tumors yielded significant intratumoral bioactive C6 accumulation by the EPR effect, but tumor growth was not controlled. TA increased bioactive C6 accumulation by ∼12.5-fold over the EPR effect. In addition, TA+CNL caused shifts in long-chain to very-long-chain ceramide ratios (i.e., C16/24 and C18/C24) that could potentially contribute to tumor control. Nonetheless, these changes in intratumoral ceramide levels were still insufficient to confer tumor growth control beyond that achieved when combining with TA with control “ghost” nanoliposomes (GNL). While this lack of synergy could be due to increased “pro-tumor” sphingosine-1-phosphate (S1P) levels, this is unlikely because S1P levels exhibited only a moderate and statistically insignificant increase with TA+CNL. In vitro studies showed that 4T1 cells are highly resistant to C6, offering the most likely explanation for the inability of TA to synergize with CNL. Thus, while our results show that sparse scan TA is a powerful approach for markedly enhancing CNL delivery and generating “anti-tumor” shifts in long-chain to very-long-chain ceramide ratios, resistance of the tumor to C6 can still be a rate-limiting factor for some solid tumor types.

## Introduction

Sphingolipid metabolism is crucial in the regulation of cancer cell growth and therapeutic resistance^1,2^. One of the most important classes of sphingolipid metabolites are ceramides, which act as second lipid messengers^2^. A variety of stimuli (e.g., chemotherapy, radiation, and antibodies) can initiate ceramide-mediated cell signaling leading to cell senescence, cell cycle arrest and apoptosis^1–3^. However, a deficiency of intracellular ceramides by increased synthesis into other compounds (e.g., glucosylceramides via glucosylceramide synthase^2^) contributes to chemotherapeutic resistance^1^. Additionally, exogenous ceramide delivery in combination with chemotherapy has shown efficacy above each monotherapy in murine melanoma tumors^4^ and human pancreatic cancer cells^5^. Specifically, short chain C6-ceramide (C6), has been shown to decrease human breast cancer cell viability and decrease murine breast tumor growth when package into a nanoliposome (CNL)^6^.

With a need for targeted, non-invasive, non-ionizing therapy on the rise, focused ultrasound (FUS) offers a versatile platform from which to treat a variety of diseases, including cancer, while mitigating side effects. FUS is essentially the concentrated deposition of soundwave energy into a small volume with thermal (continuous wave) and/or mechanical effects (pulsed waveform). Thermal effects range from tissue hyperthermia to coagulative necrosis, whereas mechanical effects run the gamut from endothelial cell sonoporation to microvascular ablation. Interestingly, FUS and radiation may synergize as ultrasound-mediated microbubble (USMB) cavitation has been used for increasing cell death and decreasing tumor growth in combination with radiation therapy^7–11^. Importantly, USMB cavitation increases total ceramide signaling. This therapeutic effect is further increased when USMB cavitation is interwoven with radiation therapy in the PC3 prostate tumor model^7,9,10^.

In studies to date, CNL targeting to tumors has been done passively, taking advantage of the enhanced-permeation and retention (EPR) effect, which is created by “leaky” tumor vasculature. However, we hypothesize that a more direct targeting approach could result in more efficacious delivery and treatment. While we have published extensively on using USMB cavitation for enhanced drug and gene delivery^12–20^, with much of this work centered on nanoparticle delivery to solid tumors, the most widely used clinical FUS regimen is partial thermal ablation (TA). TA is easy to accomplish with standard electrical equipment given its relatively low-pressure and long timescale (seconds to minutes) requirements. Further, TA has been associated with antitumor immune responses in several types of cancer, including pancreatic^21^, prostate^22– 24^, colon^25^, kidney^26,27^, breast^28^ and melanoma^29^. Many other investigators have also shown that thermal-based FUS can facilitate enhanced delivery of systemically administered payloads to ultrasound-targeted tumor tissue, including when a fraction of the tumor volume is thermally ablated^30^ or undergoes hyperthermia^31–33^ (i.e., represents the peri-ablative zone of thermally ablative treatments).

Here, we tested the hypothesis that TA will augment CNL monotherapy in murine triple-negative breast cancer (TNBC). More specifically, we predicted that TA would increase intratumoral bioactive C6-ceramide concentration, thereby increasing tumor growth control above either monotherapy. Overall, our results show that, while TA does indeed yield a substantial increase in CNL/C6 ceramide delivery to 4T1 breast tumors, as well as presumably beneficial shifts in intratumoral long-to-very-long chain ceramide ratios, these changes are insufficient to enhance tumor control, presumably due to marked inherent resistance of 4T1 tumor cells to C6 ceramide-mediated tumor cell killing.

## Materials and methods

### Cell line and animal maintenance

The 4T1 cell line was maintained in RPMI-1640+L-Glutamine (Gibco #11875-093) supplemented with 10% Fetal Bovine Serum (FBS, Gibco #16000-044) at 37°C and 5% CO2. Thawed cells were cultured for up to three passages and maintained in logarithmic growth phase for all experiments. Cells tested negative for mycoplasma prior to freezing.

All mouse experiments were conducted in accordance with the guidelines and regulations of the University of Virginia and approved by the University of Virginia Animal Care and Use Committee. Eight-week-old to ten-week-old female Balb/c mice were obtained from NCI Charles River (NCI CRL #028) and The Jackson Laboratory (Jax #000651). 4×10^5^ 4T1 cells diluted in ice-cold 1X DPBS (Gibco, #14190-144) were subcutaneously (s.c.) implanted into the right flank of mice after shaving through a 25G x 1 ½ in needle (BD PrecisionGlide Needle #305127). Mice were housed on a 12-hour/12-hour light/dark cycle and supplied food ad libitum. Tumor outgrowth was monitored via digital caliper measurements. Tumor volume was calculated as follows: volume = (length×width^2^)/2. Fourteen days following tumor implantation, mice were randomized into groups in a manner that ensured matching of mean starting tumor volume across experimental groups.

### In vivo ultrasound-guided partial thermal ablation

Mice underwent Sham or TA treatment 14 days post-inoculation. On treatment day, mice were anesthetized with an intraperitoneal (i.p.) injection of ketamine (50 mg/kg; Zoetis) and dexdomitor (0.25 mg/ kg; Pfizer) in sterilized 0.9% saline (Hospira #PAA128035). Dexdomitor was reversed with a s.c. injection of atipamezole hydrochloride (0.25 mL in 10 mL sterilized 0.9% saline, 0.4 mL s.c., Antisedan, Zoetis) after Sham or TA treatment. Right flanks of mice were shaved, after which TA was performed using an in-house built ultrasound-guided FUS system. This includes incorporation of ultrasound visualization/guidance orthogonal to the focal axis of the therapy transducer. The system uses a linear imaging array (Acuson Sequoia 512, 15L8 imaging probe, 8 MHz, 25 mm field width) and a 1.1 MHz center-frequency, single-element therapy transducer [H-101 (Sonic Concepts Inc., Bothel, WA). The therapy transducer had an active diameter of 64 mm and radius of curvature of 63.2 mm. The transducer was operated at third harmonic (3.28 MHz), with a -6dB focal size of 0.46 mm x 0.46 mm x 3.52 mm = ∼0.39 mm^3^. Both the imaging and treatment transducers were ultrasonically coupled using degassed, deionized water at 37°C. TA was applied continuously for 10 s, at a peak negative pressure = 12 MPa, with treatment points spaced 1 mm in a rectangular grid pattern and 2 planes of treatment, which were separated by 2 mm. The treatment scheme is outlined in Figure S1. Sham treatment comprised of partially submerging the mice in the 37°C water bath for 6 minutes.

### Nanoliposome delivery

The formulation of C6-ceramide nanoliposomes (CNL) was done as previously described^34^. Control “Ghost” nanoliposomes (GNL) were formulated with the same lipid composition, without C6-ceramide loading. Both CNLs and GNLs were intravenously (i.v.) injected at 36 mg/kg mouse. The timing of the i.v. injection relative to Sham or TA treatment was immediately before, except for one study wherein they were injected either 1h or 24h before TA or Sham. For tumor growth control studies, CNL or GNL injections started on day 12 and occurred every two days until day 20 post-inoculation.

### Tumor and serum collection

Mice were euthanized with Euthasol® (Virbac, Inc., 390 mg pentobarbital sodium and 50 mg phenytoin sodium per mL, mice were given 0.05 mL of stock i.p.). Tumors and blood were collected at times specified in figures. Tumors were resected with adjoining non-tumor tissue removed and stored at -80°C in 1X DPBS (Gibco, #14190-144). Blood was collected (BD, Microtainer SST #365967) via cardiac puncture and allowed to clot for 30 minutes as per the manufacture’s instructions. The clotted blood was centrifuged (Eppendorf, #5424) for 90 s at 15000 RPM to separate whole blood from serum. 60 μL of serum was placed in a microcentrifuge tube (Fisherbrand, #02-681-343) and stored at -80°C.

### Mass spectrometry

Lipid extraction and analysis was done using liquid chromatography-electrospray ionization-tandem mass spectrometry (LC-ESI-MS/MS). Lipids were extracted from tumors/serum using an azeotrophic mix of isopropanol:water:ethyl acetate (3:1:6; v:v:v). Internal standards (10 pmol of d17 long-chain bases and C12 acylated sphingolipids) were added to samples at the onset of the extraction procedure. Extracts were separated on a Waters I-class Acquity UPLC chromatography system. Mobile phases were (A) 60:40 water:acetonitrile and (B) 90:10 isopropanol:methanol with both mobile phases containing 5 mM ammonium formate and 0.1% formic acid. A Waters C18 CSH 2.1 mm ID × 10 cm column maintained at 65°C was used for the separation of the sphingoid bases, 1-phosphates, and acylated sphingolipids. The eluate was analyzed with an inline Waters TQ-S mass spectrometer using multiple reaction monitoring. All data reported are represented as pg/mg protein/g tumor unless specified otherwise.

### In-vitro cell viability assays

4T1 cells were seeded on 96 well plates. After 24 hours, cells were treated with CNL or ghost liposomes at the indicated concentrations. MTS assays were performed according to the manufacturer’s instructions (Promega, Madison, WI). Absorbance at 490nm was determined with a Cytation 3 plate reader (Bio Tek, Winooski, VT).

### Statistical Analysis

All statistical analyses were performed in GraphPad Prism 9 (GraphPad Software). Tumor growth comparisons between treatment groups were performed with a repeated-measures mixed-effects model with three factors (i.e., Time [repeated-measures], GNL/CNL, and Sham/TA) and corresponding interaction terms using the Geisser-Greenhouse correction. Assuming normal distributions, outliers were identified with the ROUT method with the strictest tolerance (false discovery rate or Q=0.1%) to remove more definite outliers from mass spectrometry data. For the mass spectrometry data (e.g., C6-ceramide levels), a one-way Brown-Forsythe and Welch analysis of variance (ANOVA) and Dunnett T3 post-hoc tests were performed with the one-factor being injection groups (i.e., No Injection, GNL, and CNL). When comparing the three injection groups and Sham/TA treatment groups (i.e., second factor), a full-model, two-way ANOVA was performed. When comparing two groups, an unpaired, two-tailed t-test with Welch’s correction (i.e., did not assume equal standard deviations) was performed. For in vitro viability assays, non-linear least squares regression, with reported adjusted R-squared values, was performed fitting the data to the absolute IC50 equation^35,36^; the relative half maximal inhibitory concentration (IC50) was derived from said regression. All figures show the mean ± standard error of the mean (SEM). P-values and significance are specified in figure legends.

## Results

### CNL Monotherapy Enhances Intratumor C6 Ceramide but Does Not Control 4T1 Tumors

We first asked whether i.v. administration of CNLs elicits a significant increase in intratumor C6 ceramide levels in 4T1 tumors via the EPR effect. To this end, using mass spectrometry, we measured a ∼ 5-fold increase over background in bioactive C6-ceramide levels in CNL-treated 4T1 tumors when compared to GNL-treated control tumors at 24 hours post-injection. Given this result, we then tested whether a clinically-aligned CNL dosing regimen leads to 4T1 tumor control. CNLs were i.v. administered to tumor-bearing mice every other day, starting on day 12 post-inoculation. Despite the ability of CNL monotherapy to increase intratumoral C6 levels, 4T1 tumor growth was not controlled (Fig. 1b).

**Fig. 1.**
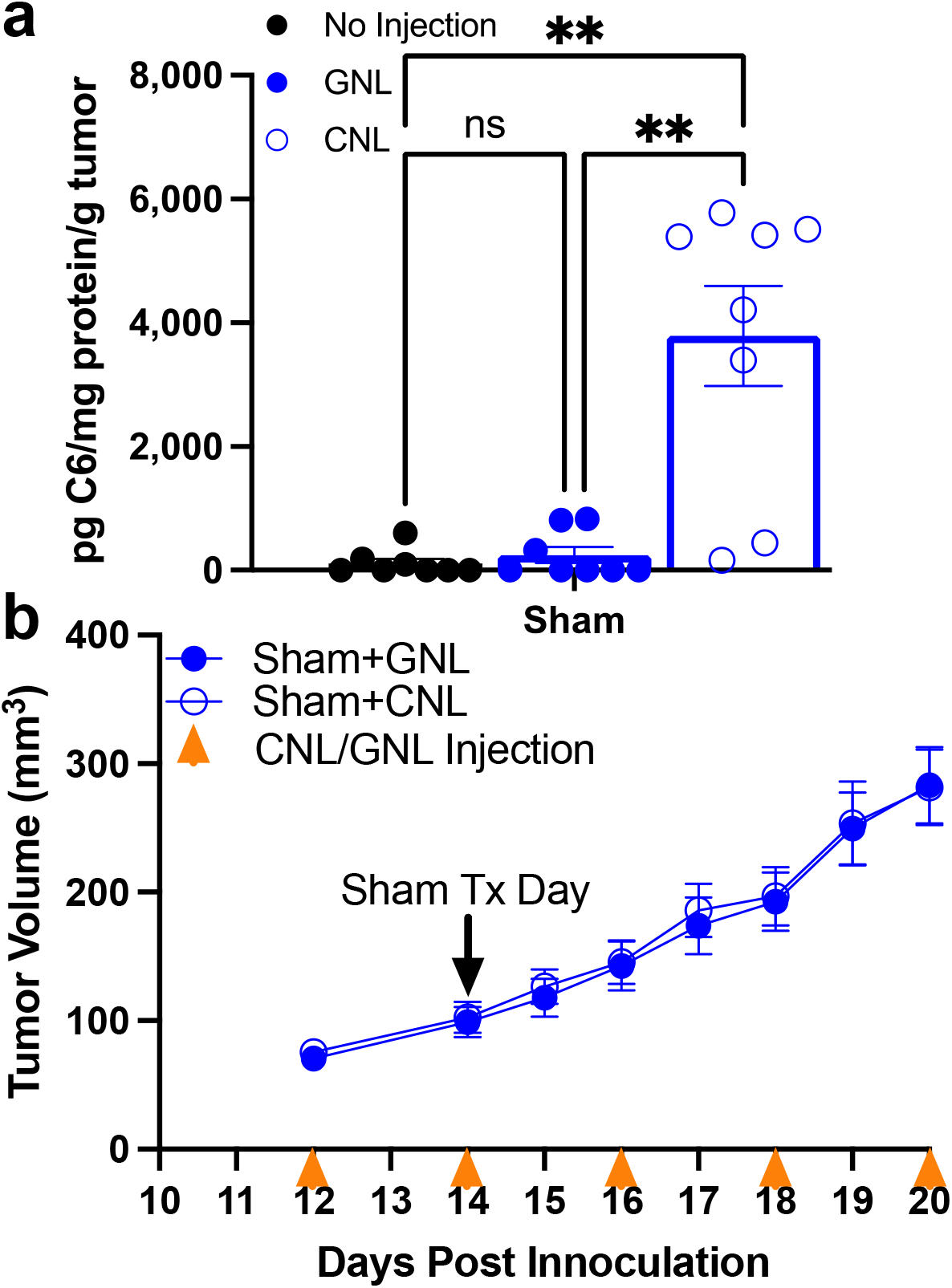
4T1 tumors are resistant to CNL monotherapy. a. C6-ceramide levels in 4T1 tumors 24 hrs post Sham treatment (n=8-9) (*p<0.05). b. Tumor growth curves indicate that CNL monotherapy does not control 4T1 tumor growth (n=14).

### Spare Scan FUS Thermal Ablation Further Augments Intratumoral Bioactive C6-Ceramide Accumulation

Given considerable evidence that TA of solid tumors with focused ultrasound can enhance liposome delivery to tumors^30,37^, we tested whether such an approach could augment CNL delivery to 4T1 tumors, above that achieved by the EPR effect. To this end, we combined CNL administration with the TA scheme outlined in Fig. S1. A preliminary study was first performed to establish which relative timing between CNL administration and TA yields maximum C6 ceramide levels in tumors. We looked specifically at cases wherein CNLs were injected at the time of TA (i.e., 0 h before FUS), 1h before TA, or 24h before TA. As shown in Fig. 2a, injecting CNL at the time of TA yielded the highest intratumor C6 ceramide levels. Thus, for all remaining experiments, CNLs were injected at the time TA was applied. Next, we compared intratumoral C6 levels generated by TA+CNL to those generated by CNL injection alone (i.e., the EPR effect). Notably, TA conferred an ∼12.5-fold increase in intratumor C6 ceramide above the EPR effect (Fig. 2b), indicating that TA markedly augments CNL delivery to solid tumors.

**Fig. 2.**
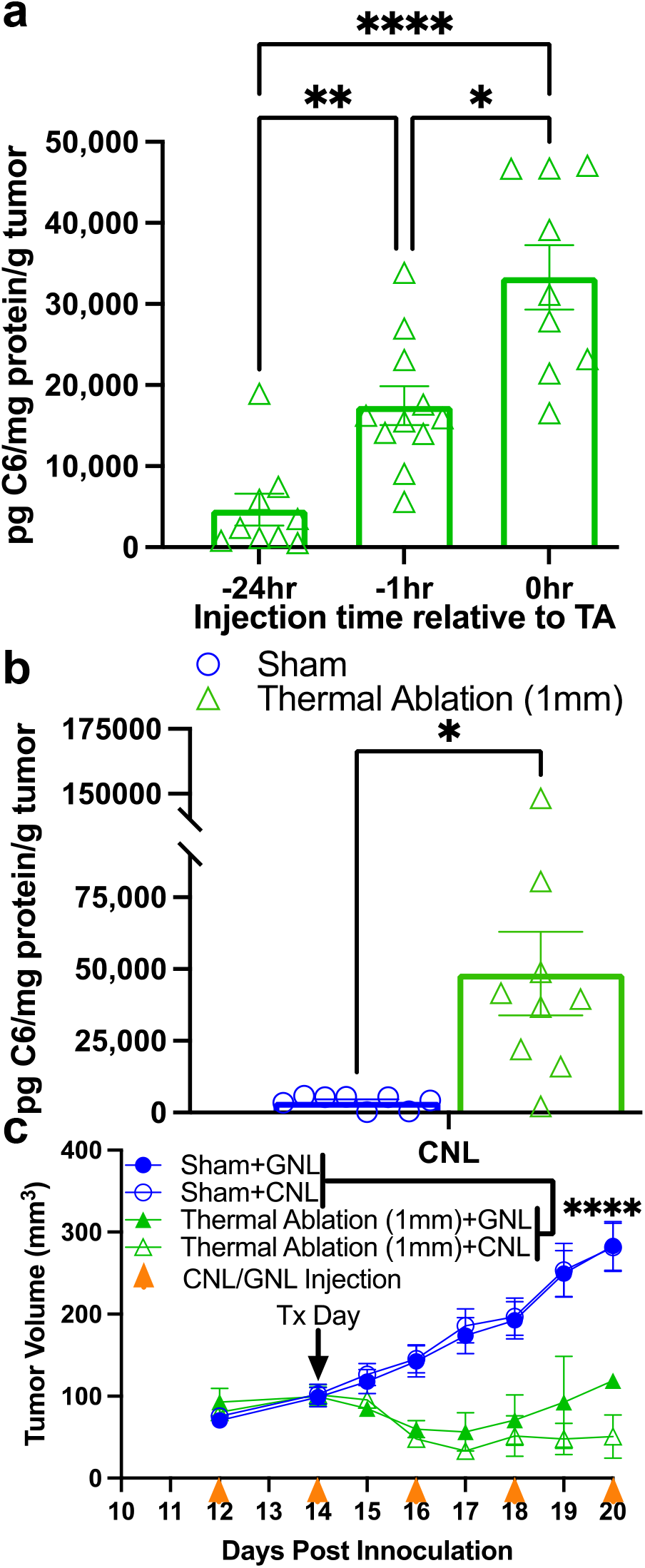
TA markedly increases C6 ceramide levels but does not synergize with CNL in controlling 4T1 tumor growth. a. C6-ceramide levels 24 hrs after TA with three different CNL injection timings relative to TA (n=9-11) (*p<0.05, **p<0.01, ****p<0.0001). b. C6-ceramide levels from Sham and TA treated tumors 24 hrs post treatment (n=8-9) (*p<0.05). CNL were injected at the time of TA or Sham treatment. Results confirm that injecting CNL at the time of TA is optimal for C6 delivery. c. Tumor growth curves indicate that, while TA controls 4T1 tumor growth, no further statistically significant benefit is conferred by CNL (n=14-16) (****p<0.0001).

### CNL Administration Does Not Further Augment 4T1 Tumor Control with FUS Thermal Ablation

After establishing the ability of TA to augment CNL delivery, we tested whether CNL administration would cooperate with TA in controlling 4T1 tumors. CNL and GNL were injected every other day beginning on Day 12 post-inoculation. TA or Sham treatment was applied on Day 14, and tumor volumes were measured every other day. Here, when combined with both CNL and GNL injection, TA robustly controlled 4T1 tumor growth when compared to the Sham treated groups (Fig. 2c). However, despite yielding enhanced intratumor C6 ceramide levels, CNL administration in combination with TA did not improve tumor control beyond that achieved when TA was combined with administration of control GNLs (Fig 2c).

### Modulation of Ceramide Metabolite Levels in Tumors Treated with TA and CNLs

We next sought to generate hypotheses for why TA was not cooperating with CNLs in the control of 4T1 tumor growth. One possible explanation was that combining CNLs with TA was shifting other (i.e., not C6) ceramide metabolite levels such that tumor control synergy was lost. It is, for example, known that higher levels of shingosine-1-phosphate inhibit tumor control^38–40^. Interestingly, as shown in Fig 3a, the ratio of C16 to C24 ceramide trended toward an increase when CNL was combined with TA. Furthermore, combining TA with CNL led to a statistically significant doubling of the ratio of C18 to C24 ceramide. In essence, when examining the underlying absolute numbers, long-chain ceramide (i.e., C14, C16, C18) levels tended to increase with TA+CNL, while very-long-chain ceramide (i.e., C24, C26) levels remained unchanged (Fig S3). Thus, it is unlikely that changes in long and very-long chain ceramide levels account for the lack of synergy between TA and CNL. Indeed, these levels are actually shifted to a presumably more beneficial anti-tumor state when TA is combined with CNLs. When examining S1P levels, we did observe a trend toward an increase with TA+CNL (Fig 3a). Although not statistically significant, it is possible that S1P levels could have contributed to a lack of tumor growth control synergy between TA and CNL.

**Fig. 3.**
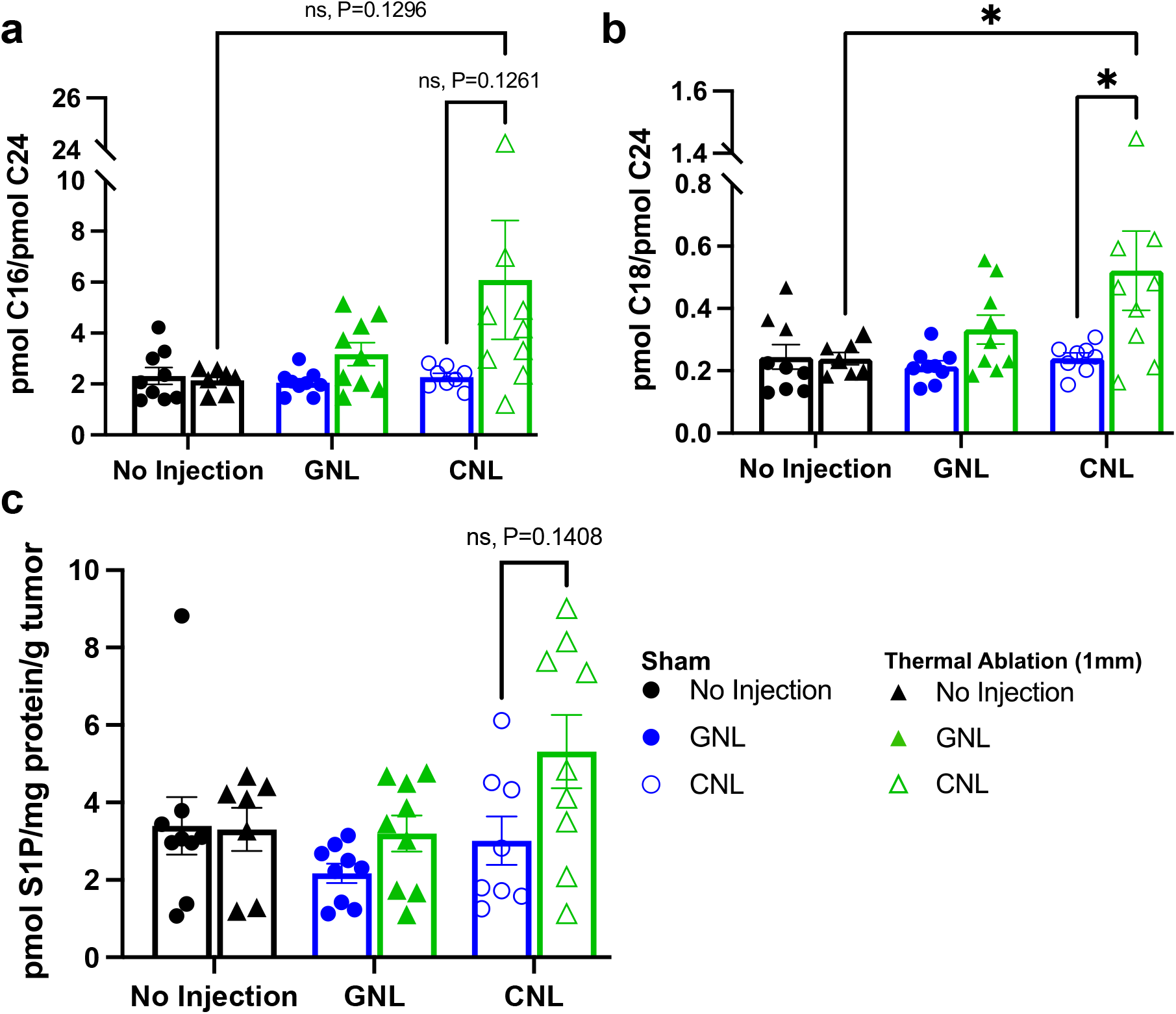
Ceramide metabolite levels in 4T1 tumors measured 24 hrs after TA or Sham treatment in combination with GNL or CNL. a. Ratios of C16 long-chain to C24 very-long-chain ceramide. b. Ratios of C18 long-chain to C24 very-long-chain ceramide. c. Sphingosine-1-phosphate (S1P) levels. For all data sets: n=7-9 and *p<0.05.

### 4T1 Cells Are Inherently Resistant to CNL Therapy

Finally, we tested whether the lack of synergy between TA and CNL could be caused by 4T1 cells possessing an unusually strong inherent resistance to CNL therapy. To this end, cell viability assays were run for 4T1 cells exposed to varying concentrations of CNLs in-vitro for either 24h or 48h. Here, we found that IC50 values for 4T1 cells exposed to CNLs at 24hrs and 48hrs were 33.83 and 17.17 μM, respectively. These values were obtained based on the non-linear least square fit of the data (Fig. 4).

**Fig. 4.**
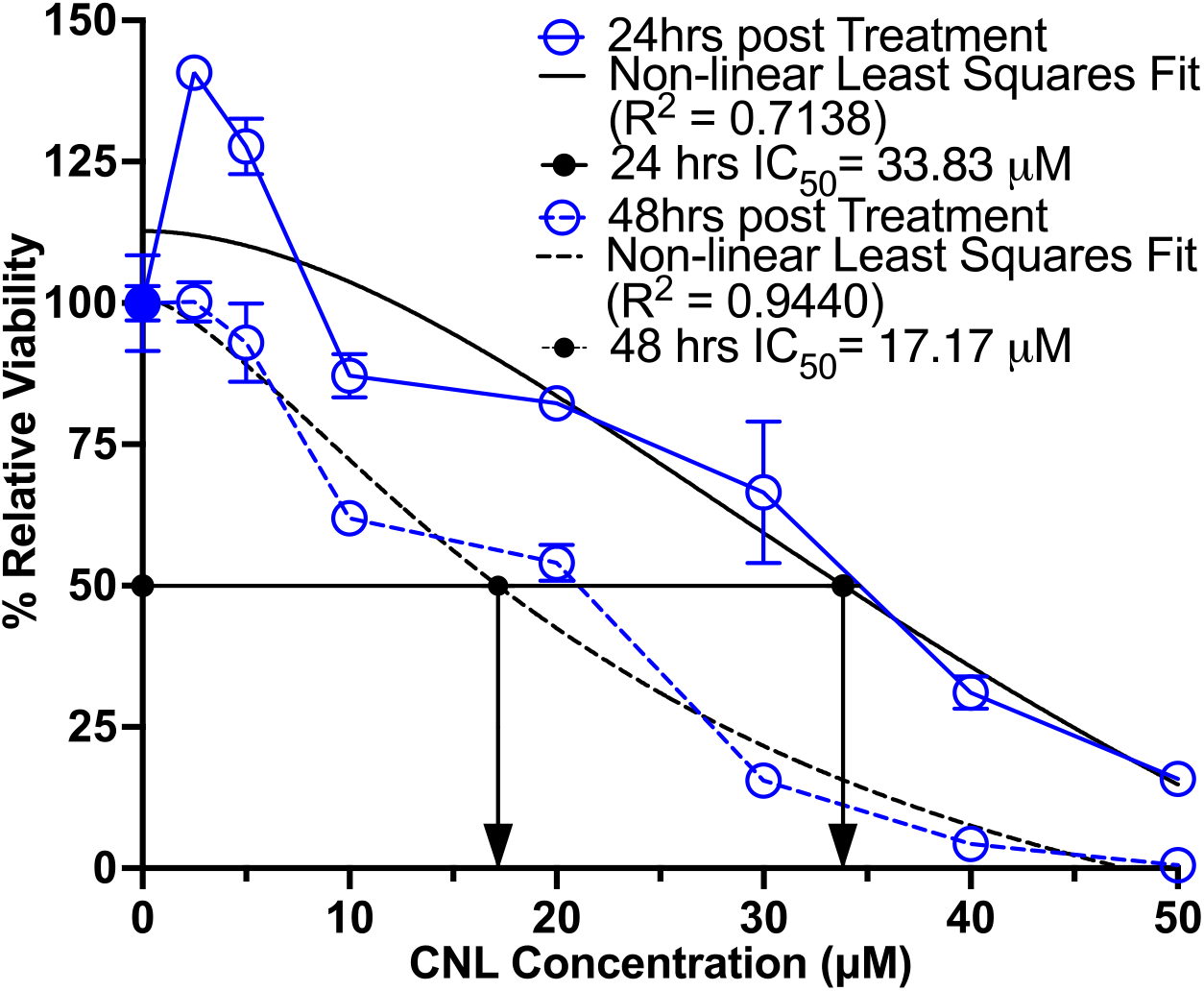
4T1 tumor cells are resistant to CNL monotherapy. *In vitro* viability of 4T1 cells exposed to various concentrations of CNL for either 24 or 48 hours (n=3-4). Viability is relative to GNL control group (n=10). Data was fitted with non-linear least square fit (black lines) from which the relative IC50 was calculated (black arrows).

## Discussion

The overall goal of this study was to determine whether sparse scan TA cooperates with intravenous CNL administration to better control solid tumor growth. We first established that, while the EPR effect permits the delivery of C6-bearing CNLs to the 4T1 TNBC microenvironment, tumor growth was not controlled. Knowing that TA increases intratumoral nanoparticle delivery from other studies^28,30,41,42^, we next tested whether TA increases intratumoral C6 as well. Even though we were able to significantly increase intratumoral C6 concentrations, the combination of TA and CNLs proved no more effective in controlling tumor growth than TA alone. Since an increase in ceramide content is considered pro-apoptotic^38,43,44^ while conversion of ceramides into S1P by sphingosine kinases is pro-tumorogenic^39,45^, we then investigated a variety of intratumoral ceramides of different chain lengths and S1P concentrations. We found that long-chained ceramide concentrations were significantly increased when treating tumors with CNLs and TA; S1P was only trending in its increase. Given the increased CNL delivery and the skewing of the ceramide metabolite landscape to anti-tumor, we tested whether 4T1 tumor cells were inherently resistant to the CNL therapy and found them to indeed be unusually resistant. We conclude that TA markedly enhances CNL delivery to 4T1 breast tumors and favorably shifts ceramide metabolism; however, these beneficial changes are insufficient to achieve synergistic tumor control, presumably due to exceptionally high resistance of 4T1 cells to C6 ceramide.

Our present findings indicate that sparse scan TA induces a substantial (∼12.5 fold) increase in ceramide delivery over the EPR effect, which is appreciably above that reported by others combining TA and hyperthermia with other liposomal delivery platforms. For example, hyperthermia has been shown to (i) increase intratumoral liposomal delivery by 50-80% in Met-1^fvb-2^ mammary fat pad tumors 18 hours after treatment^31^, (ii) increase doxorubicin (DOX) fluorescence (i.e., surrogate for DOX delivery) by ∼ 3.4 fold in NDL mammary carcinoma post-hyperthermia^46^, and (iii) increase liposomal DOX delivery to murine mammary adenocarcinoma (JC cell line) increased by ∼ 3 and 0.5 fold 5 min and 120 minutes after hyperthermia, respectively, over liposome alone^47^. In studies that combined TA with liposomal delivery, ^111^In-labeled liposome delivery increased from TA treatment by ∼ 120% after 90 minutes and ∼ 33% after 48hrs compared to sham in hindleg rhabdomyosarcoma, while DOX delivery increased by ∼ 3 fold 90 minutes after injection^37^. Furthermore, ^64^Cu PET labeled liposomal delivery increased by 3-4 fold compared to contralateral tumors in NDL mammary carcinoma 3, 20 and 48 hours post TA, while, in the same study, liposomal accumulation in 4T1 tumors increased by ∼ 2 fold^30^. Interestingly, 4T1 has been shown to allow more (∼ 2 fold) passive intratumoral liposome accumulation than the NDL breast cancer model^30^, which indicates a higher EPR effect in the 4T1 model. However, in our subcutaneous 4T1 breast cancer model, we saw a ∼ 12.5-fold increase in intratumoral bioactive ceramide after TA treatment. One possible explanation as to why our TA regimen increased relative ceramide levels by such a high degree compared to that of the aforementioned study with NDL and 4T1 tumor models is that each sonication in a grid treatment pattern was immediately sequential, while in Wong et. al. performed their grid TA regimen with a 30 second wait period between sonications^30^.

Because changes in endogenous ceramide levels (i.e., not C6) elicited by TA could also synergize with, or compete with, C6 in tumor control, we measured long-chain and very-long-chain ceramides after treatment. When combining our TA regimen with CNL, we found the ratio of C16 to C24 ceramide increased, while the ratio of C18 to C24 ceramide doubled. The corresponding absolute numbers of long-chain ceramide (i.e., C14, C16, C18) levels tended to increase with our combination therapy, while very-long-chain ceramide (i.e., C24, C26) levels remained stable. Though these results are compelling, they are difficult to interpret, as there is considerable contradiction in the literature regarding endogenous ceramide levels and tumor control. For instance, in human colon cancer cell lines, C2, C6 and C18 therapies inhibited cell growth via the cytochrome release pathway while normal liver cells remained resistant^48^. In head and neck squamous cell carcinomas, C18 was downregulated and alleviation of this dysfunction by restoring intracellular C18 concentrations decreased cell growth^49^. However, contrary to what was found in the human colon cancer cell lines, knockdown of C16 was proapoptotic while the restoration thereof was prosurvival^50^. Human breast cancer tissue and cell lines exhibit increased intracellular production of C16, C24 and C24:1^51^. Overexpression of the ceramide synthases (CerS) 4 and 6 (i.e., CerS that metabolize C14, C16, C18 and C20) led to apoptosis and cell proliferation inhibition^51^, while overexpression of CerS 2 (i.e., CerS that metabolize C22 and C24) led to increased cell proliferation^51^, chemosensitivity^52^, and inhibited migration and invasion^53^. With this seeming contradicting data, strong conclusions about our findings regarding ratios and absolute ceramide concentrations are hard to draw. Going forward, studies specifically designed to unravel these putative mechanisms would likely be required to determine which ceramides and CerS may be contributing to CNL and TA resistance in the 4T1 model and breast cancer at large. Our primary hypothesis for why markedly enhanced C6 delivery with TA did not further control 4T1 tumors is that this cell line is unusually resistant to C6. Indeed, we determined that IC50 values for 4T1 cells exposed to CNLs at 24hrs and 48hrs were 33.83 and 17.17 μM, respectively. In comparison, in a previous study^6^, Stover et al. showed that, in the CNL-responsive 401.4 breast tumor model, the IC50 at 48h of CNL exposure was 5 μM, which is ∼3.5-fold lower than the value for 4T1 cells obtained here. Furthermore, previous studies on prostate cancer cell lines (i.e., PC-3, PPC-1 and DU145) that respond to CNLs have yielded IC50 values of ∼10-12 μM^54^. Thus, it is likely that, despite the ability of TA+CNL to markedly enhance C6 ceramide levels in 4T1 solid tumors, further 4T1 tumor growth control is not realized due to inherent resistance to C6 ceramide.

Though this particular cancer model was resistant to CNL therapy, this study further establishes that sparse scan TA, as applied here, is a readily available and clinically translational option for substantially enhancing the delivery of the CNL formulation. Since TA can increase therapeutic index via increased local delivery of systemically administered therapeutics, this platform holds promise in that lower systemic doses can be administered but still achieve therapeutic efficacy. Given the variety of possible cellular responses to ceramide-based therapies, this study suggests that investigation into which cancer indications are well-suited to ceramide-based therapies is vitally important.

## Supporting information

Supplemental Inofrmation

## Acknowledgments

Supported by National Institutes of Health Grants NIH R21CA230088 to M.K. and R.J.P. and R01EB030007 to R.J.P.

## Author Contributions

Conceptualization: E.A.T., M.K., R.J.P.; Methodology: E.A.T., M.K., R.J.P.; Formal analysis and investigation: E.A.T., T.F., T.D., L.R.V, N.D.S.; Writing - original draft preparation: E.A.T.; Writing - review and editing: E.A.T., T.D., T.F., N.D.S., R.J.P.; Funding acquisition: M.K., R.J.P.; Resources: M.K., R.J.P.; Supervision: M.K., R.J.P.

## Notes

### Competing Interest Statement

The authors have declared no competing interest.

