## Supplemental Inofrmation for "Solid Tumor Treatment via Augmentation of Bioactive C6 Ceramide Levels with Thermally Ablative Focused Ultrasound"

*Corresponding Author:

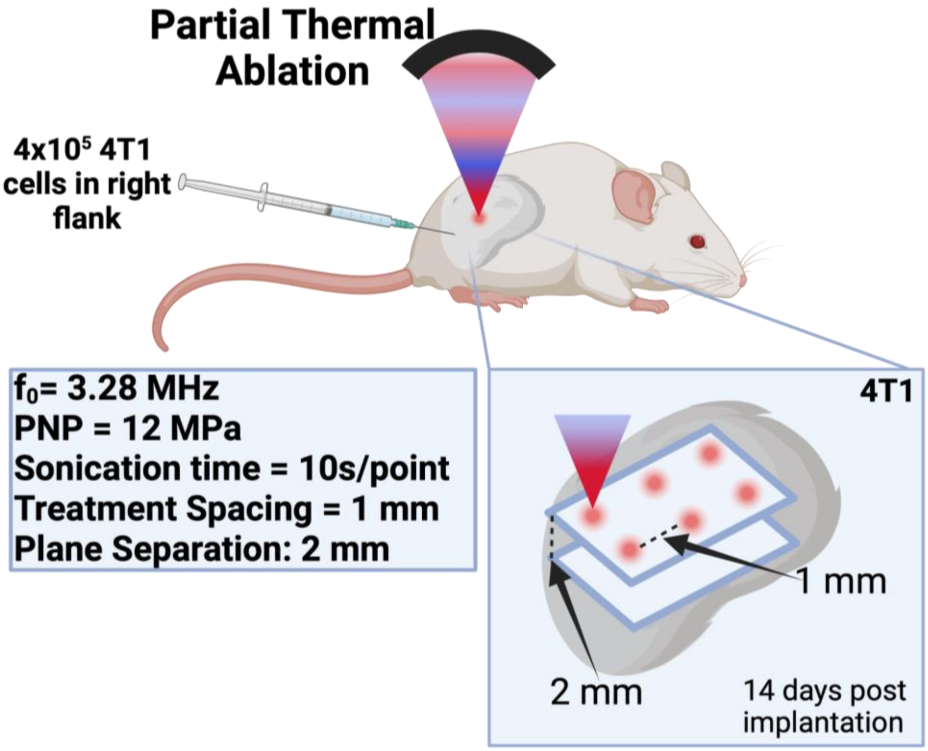


**Fig. S1 Treatment scheme for TA of 4T1 tumors.** Biorender.com

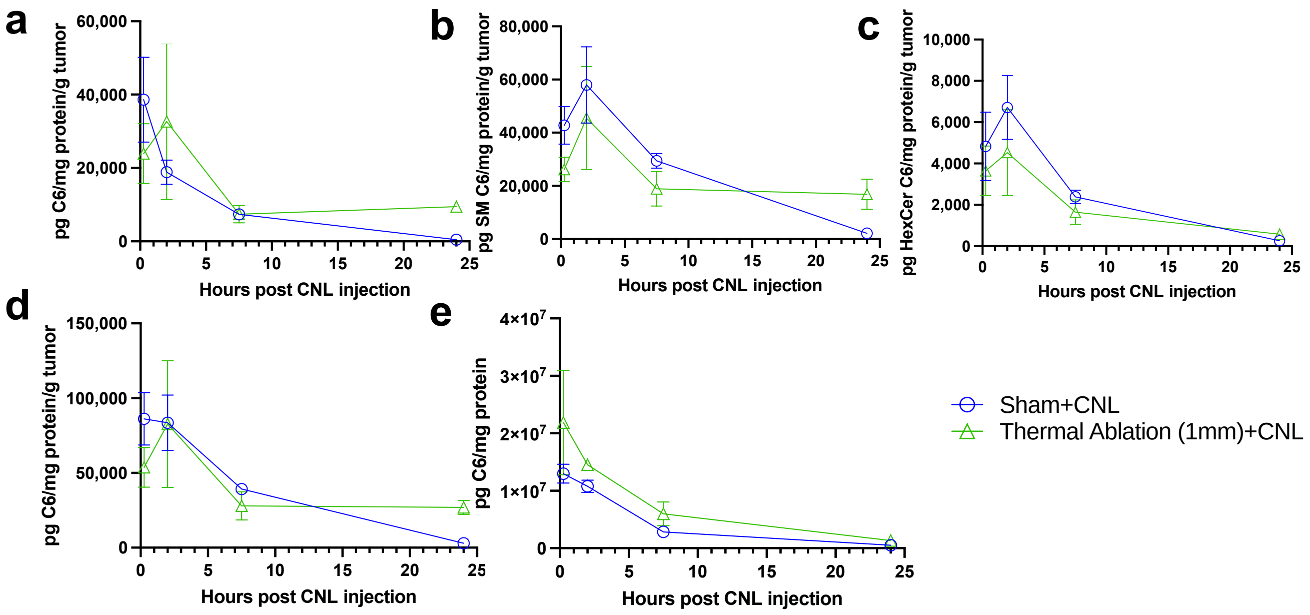


**Fig. S2 C6-ceramide levels through time after TA+CNL treatment. CNLs were injected immediately prior to Sham or TA treatment.** Tumors and serum were harvested 15 minutes (n=5-6), 2 hrs (n=4), 7.5 hrs (n=4-6) and 24 hrs (n=3-5) post treatment. a. Bioactive C6-ceramide levels. b. Levels of C6-ceramide metabolized into sphingomyelin. c. Levels of C6-ceramide metabolized into hexosyl-ceramides. d. Total C6-ceramide levels. e. Bioactive C6-ceramide levels in serum.

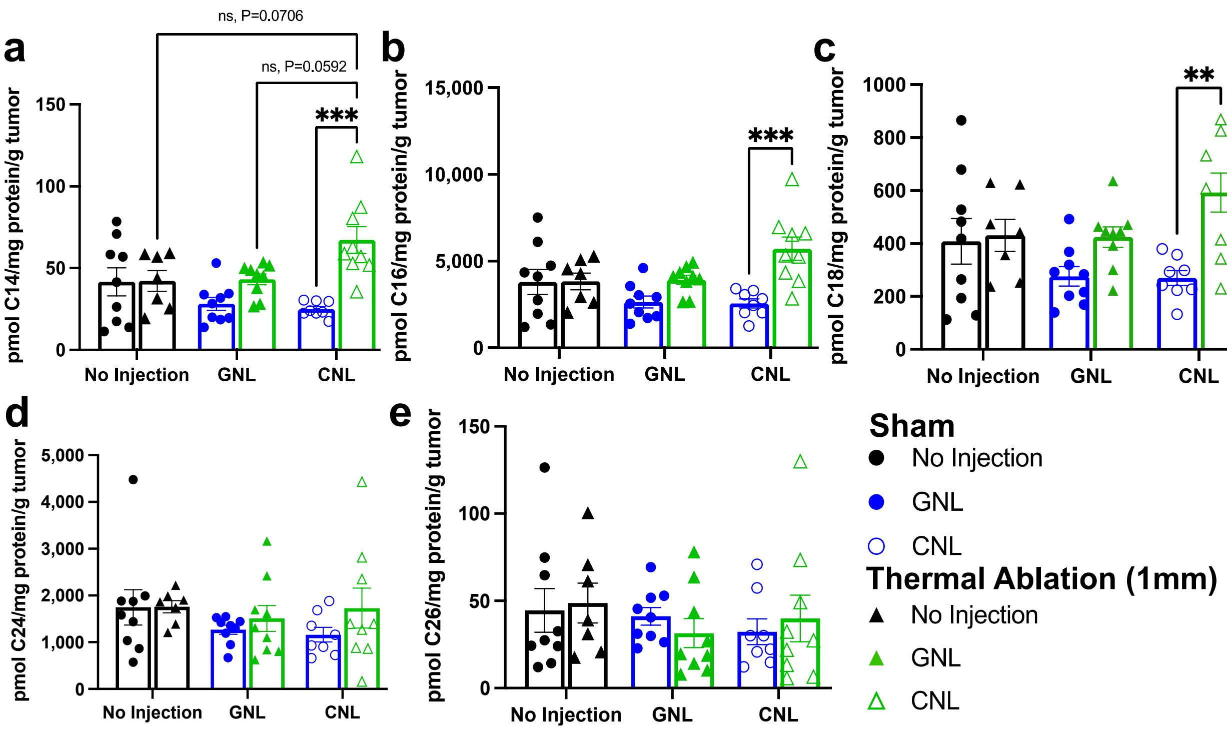


**Fig. S3 Absolute ceramide metabolite levels 24 hrs after Sham or TA treatment in combination with GNL and CNL injection.** Long-chain ceramides (A-C) are C14, C16 and C18. Very-long-chain ceramides (D-E) are C24 and C26. For all data sets: n=7-9 and **$p<0.01$, ***$p<0.001$.
